# Efficient assessment of nocturnal flying insect communities by combining automatic light traps and DNA metabarcoding

**DOI:** 10.1101/2020.04.19.048918

**Authors:** Vanessa A. Mata, Sónia Ferreira, Rebecca Campos, Luís P. da Silva, Joana Veríssimo, Martin F. V. Corley, Pedro Beja

## Abstract

1. Increasing evidence for global insect declines is prompting a renewed interest in the survey of whole insect communities. DNA metabarcoding can contribute to assessing diverse insect communities over a range of spatial and temporal scales, but efforts are still needed to optimise and standardise procedures, from field sampling, through laboratory analysis, to bioinformatic processing.
2. Here we describe and test a methodological pipeline for surveying nocturnal flying insects, combining a customised automatic light trap and DNA metabarcoding. We optimised laboratory procedures and then tested the methodological pipeline using 12 field samples collected in northern Portugal in 2017. We focused on Lepidoptera to compare metabarcoding results with those from morphological identification, using three types of bulks produced from each sample (individuals, legs and the unsorted mixture).
3. The customised trap was highly efficient at collecting nocturnal flying insects, allowing a small team to operate several traps per night, and a fast field processing of samples for subsequent metabarcoding with low contamination risks. Morphological processing yielded 871 identifiable individuals of 102 Lepidoptera species. Metabarcoding detected a total of 528 taxa, most of which were Lepidoptera (31.1%), Diptera (26.1%) and Coleoptera (14.7%). There was a reasonably high matching in community composition between morphology and metabarcoding when considering the ‘individuals’ and ‘legs’ bulk samples, with few errors mostly associated with morphological misidentification of small microlepidoptera. Regarding the ‘mixture’ bulk sample, metabarcoding identified nearly four times more Lepidoptera species than morphological examination.
4. Our study provides a methodological metabarcoding pipeline that can be used in standardised surveys of nocturnal flying insects, showing that it can overcome limitations and potential shortcomings of traditional methods based on morphological identification. Our approach efficiently collects highly diverse taxonomic groups such as nocturnal Lepidoptera that are poorly represented when using Malaise traps and other widely used field methods. To enhance the potential of this pipeline in ecological studies, efforts are needed to test its effectiveness and potential biases across habitat types and to extend the DNA barcode databases for important groups such as Diptera.

## Introduction

Recent studies have shown precipitous declines in insect populations, which can have far-reaching consequences for ecosystem functioning and thus to human lives and livelihoods (Basset and Lamarre 2019; Wagner 2020). One of the striking features of this apparent decline is that it seems to be affecting entire insect communities, rather than a few species of conservation concern or particularly sensitive species groups (Hallamann et al. 2017; Bell et al. 2020; Wagner 2020). Because of this, there is an urgent need to monitor whole insect communities and to understand the main drivers of community change over a range of spatial and temporal scales (Sánchez-Bayo and Wyckhuys 2019). This task is challenging due to taxonomic impediment (sensu, e.g., Ebach et al. 2011), which makes it hard to describe the huge diversity of insect communities with conventional methods in a cost-effective way.

The advent of next-generation DNA sequencing coupled with metabarcoding approaches is revolutionising the study of diverse insect communities by overcoming taxonomic impediment, allowing the cost-effective processing of hundreds to thousands of complex mixed community samples at high taxonomic resolution (Douglas et al. 2012; Barsoum et al. 2019; Gueuning et al. 2019; Piper et al. 2019). Typically, metabarcoding studies of communities of insects and other invertebrates involve the collection of field samples using a variety of methods, and then DNA from multi-species samples is extracted, amplified using PCR and sequenced using a next-generation sequencing platform (e.g. Braukmann et al. 2019; Marquina et al. 2019; Zenker 2020). Sequencing data is used to produce a list of taxa recorded at each site, using bioinformatic pipelines and reference libraries of DNA barcodes (Marquina et al. 2019; Zenker 2020). Applications of this general approach are increasing, particularly in the case of freshwater communities (e.g., Elbrecht et al. 2017; Bush et al. 2020), where efforts are underway to develop standardised metabarcoding approaches to be used in official monitoring programs such as the European Water Framework Directive (Hering et al. 2018). The use of metabarcoding in studies of terrestrial insects has lagged behind that of freshwater communities, but the technique has already been tested, for instance, in the monitoring of wild bees (Gueuning et al. 2019), invasive species (Piper et al. 2019) and dung insects for ecotoxicological assessments (Blanckenhorn et al. 2016), among many others. Despite these advances, however, considerable efforts are still needed to develop, optimise and standardise efficient methods to collect and process insect samples for DNA metabarcoding studies and monitoring programs, as results are conditional on methodological alternatives adopted, including DNA extraction, primer sets and bioinformatic pipelines (Brandon-Mong et al. 2016; Braukman et al. 2019; Elbrecht et al. 2019).

To the best of our knowledge, DNA metabarcoding has yet to be used for describing communities of nocturnal insects, possibly due to the bias of ecologists towards studying daytime phenomena (Gaston 2019). This is regrettable, because nocturnal insects encompass about half of all insect species, and they can be negatively impacted by factors that do not operate during the day, such as light pollution (Owens et al. 2020). Furthermore, they are key components of natural and anthropogenic ecosystems, playing significant roles as, for instance, pollinators (Macgregor et al. 2015), crop pests (Aizpurua et al. 2018) and food resources for bats and other species (Sierro et al. 2001; Mata et al. 2016; Aizpurua et al. 2018). Although nocturnal insects are often captured in passive traps aimed at sampling whole insect communities, these are often biased against some taxonomic groups. For instance, Malaise traps in combination with DNA barcoding or metabarcoding are increasingly used to survey insect communities worldwide (de Waard et al. 2019), but they are mainly effective at collecting Diptera and Hymenoptera (Matthews & Matthews 1971), and thus can underrepresent major nocturnal species groups such as moths (Lepidoptera). To overcome these limitations, studying nocturnal insect communities requires targeted sampling devices, which in the case of flying species normally involve light trapping combined with flight interception (Young 2005; Häuser and Riede 2015). The variety of light traps available is very large, ranging from commercial to customised models, and from models that are operated manually to automatic models with triggers that switch them on and off at specific times (Young 2005; Häuser and Riede 2015). Typically, a light trap can collect hundreds to thousands of individuals in a single night, and so DNA metabarcoding could help speed up and increase the taxonomic resolution of sample processing (Zenker et al. 2020). However, efforts are still needed to develop and optimise traps that can be efficiently combined with metabarcoding in large scale field surveys. Methods that can avoid the need of hand-picking individual specimens, thus reducing effort and contamination risks, and the use of chemical compounds to kill or otherwise retain insects within traps, which would make the subsequent steps of DNA extraction and amplification more difficult (Dillon et al. 1996; Ballare 2019), would be particularly useful.

In this study, we describe a methodological pipeline to study nocturnal flying insects, which combines an automatic light trap device and DNA metabarcoding. Specifically, the study aims to: (i) describe the light trap and its operation; (ii) determine its capacity in sampling a high diversity of insects in short periods of time; and (iii) test whether the molecular procedures yield estimates of species richness and composition comparable to those obtained using conventional morphological identification. Overall, the study shows the value of our new approach to facilitate the sampling of highly diverse communities of nocturnal flying insects in a short time.

## Materials and Methods

### Study design

We conducted field testing of the customised light trap in July 2017, within a protected area in north-eastern Portugal (Parque Natural Regional do Vale do Tua; 41.33 N, 7.35 W). We collected a total of 12 arthropod bulk samples, by setting 6 light traps in each of two nights in two areas of cork oak woodlands, which were expected to yield high insect diversity. The light traps were spread out so that they were not visible from one trap to the other. Evaluation of metabarcoding results used the 12 field samples, involving comparisons of species richness and composition estimated through molecular procedures *versus* conventional morphotaxonomy. We focused on species of moths (Lepidoptera) because this is a taxonomically and functionally highly diverse and relatively well-known group in the country (Corley 2015), there was a highly experienced taxonomist (MFVC) available to undertake the field identifications, and a comprehensive library of DNA barcodes was already available for a large proportion of moth species occurring in the region. Comparisons between morphological and molecular results involved three different approaches to produce the bulk samples, each representing a particular study design and objectives: (i) ‘individuals’ – bulk sample for each site produced using one individual per species present in the field sample, which can be used for studies targeting a single taxonomic group, and thus where metabarcoding of the entire bulk sample is unnecessary and may eventually introduce biases; (ii) ‘legs’ – bulk sample similar to (i), but including only one leg from each individual instead of the entire specimen, which may be used when the individuals need to be preserved for other analysis or PCR inhibitors are likely to be present in digestive tract or other tissues; and (iii) ‘mixture’ – bulk sample retaining all specimens collected, without any sorting and irrespective of taxa. In our case, the approach (iii) excluded the individuals retained for (i) and (ii), and thus did not consider the rarest species (i.e., represented by a single individual in the field sample).

### Light trap design and operation

We adapted a light trap device to allow daily deployment and automatic operation of many traps by a small field team (Figure 1). Each light trap is equipped with one-meter long IP65 3528 UV LED light strip of 395-405 nm, containing 60 LEDs of 0.08 W each, folded in three sections within a 35 cm long transparent plastic tube with 3 cm diameter, and powered by a 12 Ah 12 V lead battery. UV light was used because it is more efficient than other light sources at attracting Lepidoptera (Young 2005), which was the main focus of our study. The strip is connected to a solar light sensor that automatically activates the circuit at sunset and shuts it down at sunrise, enabling the placement of several traps throughout daytime while ensuring an equal functioning time for each of them throughout the night, avoiding bias regarding flight time activity and unnecessary draining of the battery during the day. The plastic tube is installed in the centre of 3-4 acrylic plates (50 cm × 16.6 cm × 2.5 mm), to intercept flying insects, and on top of a transparent funnel made of rigid 0.75 mm PVC film with a 4 cm diameter opening that leads into a 30 L bucket. Inside the bucket there is a breathable fabric bag containing cardboard egg boxes for the insects to rest and hide. Flying insects attracted to the light collide with the acrylic plates and eventually fall through the tunnel and get trapped inside the bucket bag. Each trap is visited in early morning by the field team, usually within one hour after dawn.

**Figure 1.**
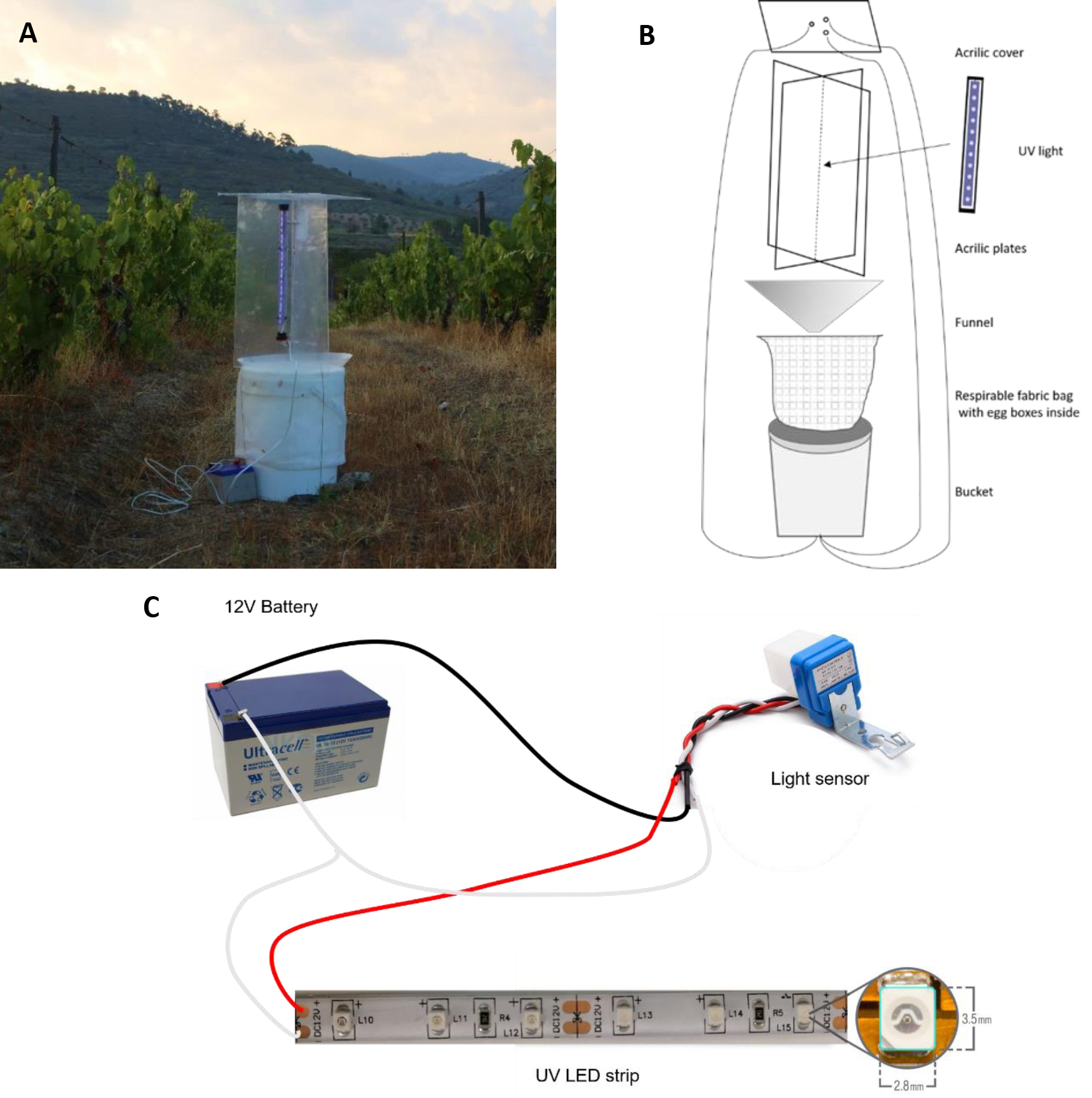
Customised UV light traps designed to collect night flying insects, showing a trap ready to operate in the field (A), a schematic representation of the assembly of all the pieces making up the complete trapping system (B), and the electric components of the trap and how to connect them (C).

### Sample processing and DNA extraction

We designed the sample processing and DNA extraction approaches to speed up processing time, reduce contamination risks and maximise DNA recovery from bulk samples. During the early-morning visit to each trap, the bag was removed from the bucket, sealed with a rubber band and frozen for at least three hours at −20 °C to immobilize the collected specimens. Each frozen sample was then thawed at room temperature. All specimens were inspected and morphological identification was attempted by a specialist (MFVC). No specimen was dissected to fully validate species identification, as the objective was to compare regular ecological studies based on simple visual identification of live moths with metabarcoding approaches. One specimen of each species identified per sample was transferred into a 50 mL falcon tube to constitute the “individual” bulk sample. From these selected specimens, a leg was removed from each and transferred into a second falcon to make up the “leg” bulk sample. Finally, all the remaining specimens (both identified and unidentified) were transferred to a third falcon to constitute the ‘mixture’ bulk sample. When very large insects (>5 cm body length) were collected, only legs were retained in the bulks to avoid biases in DNA amplification (i.e., overrepresentation of species with high biomass, and masking of rare and low biomass species). The 36 falcon tubes (three per sample) were filled with 96% ethanol and stored at room temperature until further processing. Maximum care was taken when handling specimens to avoid cross-contamination among samples, namely by thoroughly cleaning all materials used (tweezers and spatulas). Prior to DNA extraction, the ethanol was filtered out and the bulks were dried to constant weight in an incubator at 56 °C for about 2 days. Afterwards, each ‘individual’ and ‘mixture’ bulk sample was homogenized into a fine powder using the Bullet Blender 50-DX homogenizer (Next Advance, USA), with 4 glass beads of 8 mm diameter during 15 min. DNA from ‘leg’ bulk samples was extracted without further processing. DNA extraction was done using the E.Z.N.A.® Tissue DNA Kit, following an adapted protocol (Supplementary Methods). We performed up to 3 DNA extractions per sample (replicates), each one using 80 to 100 mg of the homogenized insect powder to increase species detection. In one sample (SL6) only one ‘individual’ and two ‘mixture’ replicates were possible due to low biomass.

### Metabarcoding library prep

DNA extracts were diluted 1:100 (after a small test comparing PCR amplification success at different dilutions) and amplified using the BF2-BR2 primer pair (Elbrecht and Leese 2017) in three independent reactions. PCR reactions comprised 5 µl of Qiagen Multiplex Master Mix, 0.3 µl of each 10 nM primer, 3.4 µl of H_2_O and 1 µl of diluted DNA. Cycling conditions consisted of initial denaturing at 95 °C for 15 min, followed by 35 cycles of denaturing at 95 °C for 30 s, annealing at 45 °C for 30 s and extension at 72 °C for 30 s, with final elongation at 60 °C for 10 min. The PCR products were tested in 2% agarose gel to check for the amplification success. All PCR products were diluted 1:4 with water and further subjected to a second PCR reaction in order to incorporate 7 bp long identification tags and Illumina P5 and P7 adaptors. PCR reactions were similar to that of first PCR, except that 5 uL of Phusion Hot Start Flex 2X Master Mix (NEB) was used, as well as 0.5 uL of each 10 nM indexing primer. Cycling conditions consisted of initial denaturing at 98 °C for 2 min, followed by 8 cycles of denaturing at 98 °C for 5 s, annealing at 55 °C for 20 s and extension at 72 °C for 20 s, with a final elongation at 72 °C for 1 min. PCR products were purified using Agencourt AMPure XP beads (Beckman Coulter) in a 1:0.8 ratio, quantified using Nanodrop and diluted to 15 nM. Purified and normalized PCR products were further pooled into a single library and quantified using qPCR (KAPA Library Quant Kit qPCR Mix, Bio-Rad iCycler). The final library was diluted to 4 nM and sequenced in an Illumina HiSeq 2500 Platform using a 2×250bp RapidRun kit for an average coverage of 160,000 paired reads per PCR product.

### Bioinformatic pipeline

General sequence processing was carried out using OBITools (Boyer et al. 2016), along with VSEARCH (Rognes et al. 2016) and LULU (Frøslev et al. 2017) for denoising. First, paired-end reads were aligned using the command ‘illuminapairedend’ and discarded if overlapping quality was less than 40. Second, reads were assigned to samples and primer sequences were removed using ‘ngsfilter’, allowing a total of 4 mismatches to the expected primer sequence. Finally, reads were collapsed into haplotypes using the ‘obiuniq’ command and singletons (haplotypes with only one read per sample) were removed. The remaining haplotypes left per sample were all joined in a single file and again dereplicated. The ‘--cluster_unoise’ VSEARCH command was then used to denoise the dataset by removing spurious sequences resultant from PCR and sequencing errors, followed by the command ‘- -uchime3_denovo’ to remove potential chimeric sequences. Afterwards, the remaining sequences were clustered at a 99% similarity threshold and the original reads were mapped back to the remaining haplotypes. Finally, we used LULU to remove genetically similar co-occurring haplotypes, this way highly reducing the number of mitochondrial nuclear copies present in the final dataset that tend to artificially increase the number of molecular units and taxa.

All haplotypes retained after the bioinformatic processing, were identified to the lowest possible taxonomic level, considering both moths (Lepidoptera) and other arthropod groups. The taxonomic assignment of each haplotype to a taxon was done with the support of a neighbour-joining phylogenetic tree based on an alignment of the haplotypes sorted by their read count. This allowed to visually define clusters of haplotypes that corresponded to the same taxon, as well as to identify chimeric sequences and PCR errors that remained in the dataset. Taxa were identified by comparing the representative haplotypes of each cluster against online databases (BOLD and NCBI), as well as unpublished sequences of arthropods collected in northern Portugal (InBIO Barcoding Initiative; e.g., Ferreira et al. 2020). Species level identifications were usually made for similarity values above 98.5% (da Silva et al. 2019), except in rare cases where no other species of the genus are known to exist, and thus genetic divergence reflects local haplotype diversity. Whenever a haplotype matched several species, genus, or families at similar identity levels, we selected the most inclusive taxonomic rank. For example, if a haplotype matched with 95% similarity two species of different genus belonging to the same family, we identified it only to family level. Those that best matched the same taxa were collapsed into a single taxon, as we assumed that they belonged to the same OTU. Haplotypes whose identification was only possible up to family, order or class level were clustered according to their similarity into distinct taxa (e.g., Noctuidae 1, Noctuidae 2, and so on). All identifications were checked for plausibility, considering the species geographical ranges and seasonal flight periods. Implausible identifications (e.g., species from other biogeographic regions not known to occur in the Iberian Peninsula), were reviewed and moved if needed to a higher taxonomic rank (e.g., from species to genus level).

### Statistical analysis

We compared both Lepidoptera richness and composition between morphological identification and metabarcoding approaches for each type of bulk sample. For Lepidoptera richness we applied a generalized linear mixed model (GLMM) with a gaussian distribution using the function ‘lmer’, and with light trap as random variable. Statistical significance (p < 0.05) of the model was tested with a likelihood ratio test using the function ‘anova’. To compare the species composition between the different treatments we first calculated a Jaccard distance matrix based on presence/absence records of Lepidoptera taxa in each sample, and then performed a permutational multivariate analysis of variance (PerMANOVA) using the function ‘adonis’. All statistical analyses were done in R version 3.5.2 (R Core Team 2018) using packages lme4 (Bates et al. 2015) and vegan (Oksanen et al. 2018).

## Results

### Trap operation and morphological identification

All traps operated as planned, showing their effectiveness at collecting a large number of nocturnal flying insects. Sorting of the bulk samples to extract Lepidoptera for subsequent comparisons with metabarcoding yielded a total of 871 identifiable individuals (average ± SD of 72.6 ± 23.0 individuals per sample), which were assigned to a minimum of 102 species (26.8 ± 6.7 species per sample) (Supplementary Table S1). Some specimens could not be identified visually as wing scales were largely absent and coloration patterns were no longer observable. A total of seven species accounted for 52.4% of the individuals identified, while 41 species were represented by a single individual. The individuals identified were then used to produce the bulk samples for comparison with metabarcoding. All 102 species were considered in the ‘individuals’ and ‘legs’ bulk samples, while the ‘mixture’ bulk samples included only 40 recognizable species (11.9 ± 3.6 species per sample) after removing the individuals to produce the former two bulks. However, in the ‘mixture’ bulk there were also the unidentifiable individuals, which were too damaged or otherwise lacked diagnostic features required for species level identification.

### Metabarcoding results

Considering the overall results for the three bulk types (‘individuals’, ‘legs’ and ‘mixture’), sequencing of libraries generated a total of 47,764,194 paired reads, with an average (±SD) of 151,632 ± 81,181 per PCR product. After bioinformatic filtering, we retained a total of 14,484,254 paired reads, with an average of 46,573 ± 30,371 per PCR product.

In the mixture bulk sample, we detected a total of 528 arthropod taxa (124.5 ± 33.3 taxa per sample), of which 61.2% were identified to species, 10.6% to genus, and 28.2% to family or higher taxonomic ranks. Most taxa detected belonged to Lepidoptera (31.1%), Diptera (26.1%) and 42.8% to 14 other orders (Figure 2). In the case of Lepidoptera, which were the main focus of this study, we detected a total of 189 taxa (55.2 ± 13.7 taxa per sample), of which 111 taxa (26.8 ± 8.1 taxa per sample) were represented in the ‘individuals’ bulk, 106 taxa (26.3 ± 6.7 taxa per sample) in the ‘legs’ bulk, and 164 taxa (45.8 ± 12.7 taxa per sample) in the ‘mixture’ bulk sample (Supplementary Table S1).

**Figure 2.**
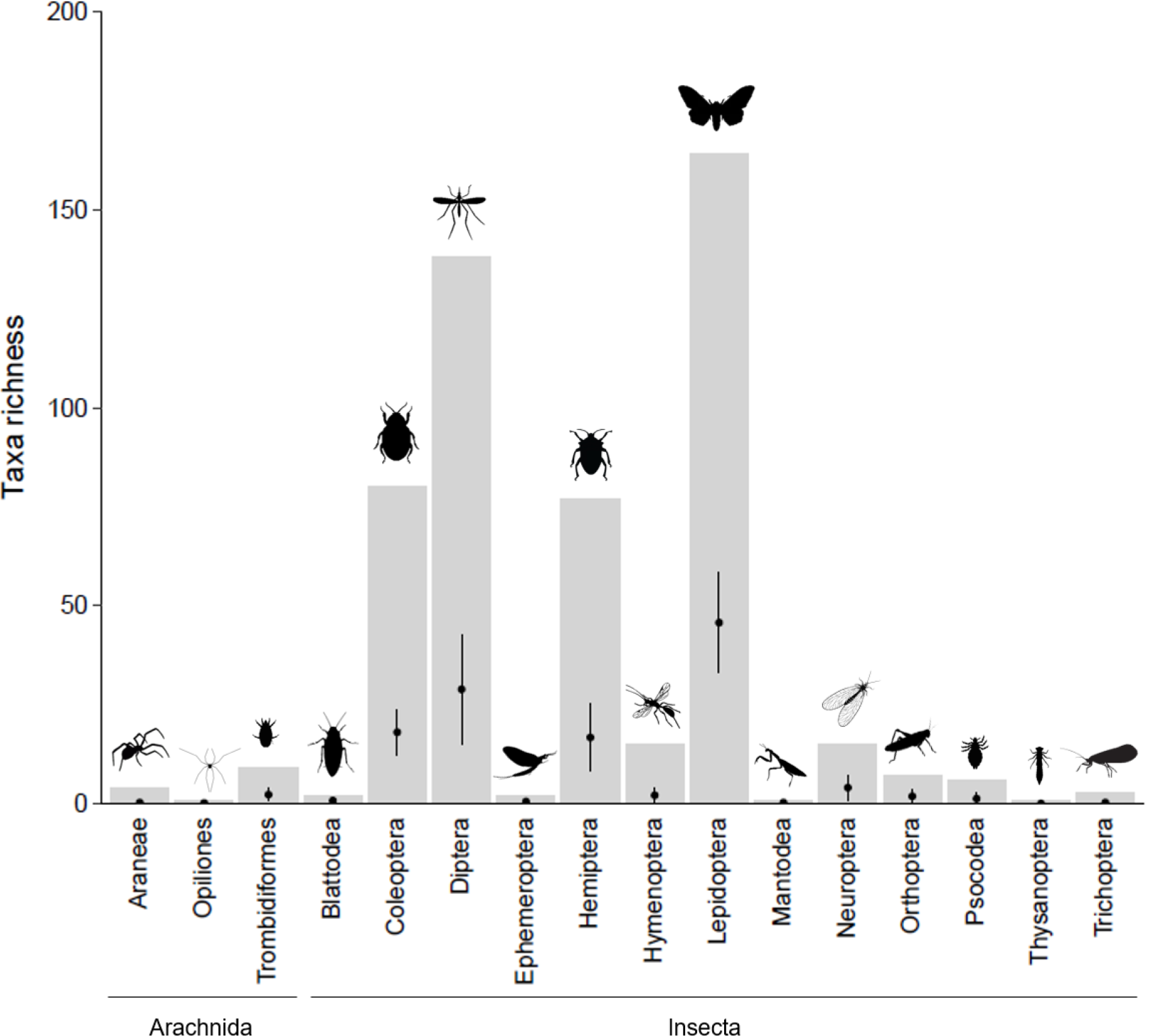
Taxa richness detected through metabarcoding in the ‘mixture’ bulk samples. Grey bars represent the total number of taxa detected per taxonomic order in the sum of all 12 samples. Whiskers represent the average ± sd number of taxa detected per sample.

### Comparisons between morphological identification and metabarcoding

Considering the Lepidoptera, comparisons between morphological and metabarcoding results were largely similar for the ‘individuals’ and ‘legs’ bulks, both in terms of species richness per sample (χ^2^ = 1.3838, df = 2, p = 0.5006) and species composition (pseudo-F = 0.8399, df = 2, R^2^ = 0.0484, p = 0.7373). The overall number of species detected was slightly higher for metabarcoding than for morphology, both for ‘individuals’ (111 *vs.* 102) and ‘legs’ (106 *vs.* 102) bulks. The mean percentage of species detected by both methods per sample was similar for the ‘individuals’ (64.6%±6.2%, 54.8-75.0%) and ‘legs’ (67.4%±6.2%, 59.4-77.4%) bulks, with similar values also for the mean percentage of species detected by morphology but not by metabarcoding (‘individuals’: 18.3%±5.0%, 11.4-29.0%; ‘legs’: 17.3%±3.6%, 11.8-21.9%), and vice versa (‘individuals’: 17.1%±5.0%, 11.1-28.1%; ‘legs’: 15.3%±3.2%, 9.7-20.0%). When considering only families normally assigned to macrolepidoptera (e.g., Erebidae, Geometridae, Noctuidae), the percentage of species shared between methods increased for both the ‘individuals’ (72.3%±10.5%, 54.5-86.4%) and ‘legs’ (74.9%±10.4%, 60.0-90.5%) bulks, while in microlepidoptera it reduced in both cases (‘individuals’: 55.4%±14.9%, 31.3-85.7%; ‘legs’: 58.7%±16.0%, 38.9-100.0%). Many of the mismatches observed were thus caused by microlepidoptera species, mainly due to species of the same genus identified differently by morphological and molecular methods, to taxa that could be identified unambiguously to species level by one method but not the other, and to more species of the same genus being detected by one of the methods. For instance, while morphology detected *Apatema mediopallidum*, metabarcoding detected only two unidentified *Apatema* species that did not match the DNA barcode of *A. mediopallidum* currently available in the databases. Likewise, morphological examination detected *Scopula submutata* and *Scythris parafuscoaenea*, while metabarcoding detected *Scopula marginepunctata* and *Scythris dissimilella*, respectively. Examples of differences in identification resolution included *Eurodachtha canigella* detected by morphology and *Eurodachtha siculella/canigella* by metabarcoding, while morphology detected an unidentified *Depressaria* while metabarcoding detected *Depressaria adustatella*. Finally, common mismatches occurred with species belonging to the genus *Idaea* and the family Brachodidae, with morphology recording *Idaea belemiata, Brachodes gaditana* and *Agonopterix scopariella*, and metabarcoding detecting *I. obsoletaria* instead *of belemiata, Brachodes gaditana* plus *B. funebris*, and *A. atomella* instead of *A. scopariella*.

In comparisons involving the ‘mixture’ bulk sample, morphology and metabarcoding recovered a significantly different community of moths. Not only the average species richness identified per sample was different (χ^2^ = 111.46, df = 1, p < 0.0001), but also the species composition (pseudo-F = 6.0464, df = 1, R^2^ = 0.2156, p = 0.001). The percentage of species detected by both methods was only 20.9%±5.5% (11.3-28.1%), with 74.6%±6.7% (64.3-83.0%) of species detected through metabarcoding but not morphology, while only 4.6%±2.1% (1.9-9.5%) of species were detected by morphology but not by metabarcoding. As for the other two bulks, the percentage of species detected by both methods was higher for macrolepidoptera (34.6%±14.6%, 10.0-58.3%) *vs.* microlepidoptera (12.8%±7.5%, 5.0-31.6%).

## Discussion

Our study provides an effective pipeline for large scale sampling of nocturnal flying insects, from field sampling using a customised trap design, through lab processing of samples, to bioinformatic analysis of sequencing data. The trap we customized proved to be successful at collecting a large number of nocturnal insects, from which bulk samples for subsequent metabarcoding processing can be easily obtained with minimal risks of contamination. We found that metabarcoding and morphological identification provide similar results when comparing bulk samples with relatively few individuals of known species, but that metabarcoding detects far more species when dealing with complex mixtures (real samples) that include many individuals that cannot be easily identified through morphological examination (e.g., due to specimen’s age, small size and scale damage, this last one probably accentuated by the transport to a freezer in a bag). Overall, we suggest that our methodological pipeline can be widely applied in ecological studies, contributing to improve our understanding of the trends and drivers of nocturnal insect communities and complement studies based on Malaise traps that are not so efficient in capturing Lepidoptera and other insects like Coleoptera, Mantodea, Neuroptera, Orthoptera and Hemiptera (Matthews & Matthews 1971).

The trap customized in our study shares many similarities to others used elsewhere (Zenke et al. 2020), including traps that are available commercially. However, it has the advantage of being relatively cheap and easy to produce, which makes it suitable for large scale field studies where resources are limited, and a large number of sites need to be sampled. Furthermore, the trap was adapted to retain insects without the need to use any chemical product to kill the individuals, which likely facilitates the subsequent steps of DNA extraction and amplification (Dillon et al. 1996; Ballare 2019). Finally, the use of a breathable fabric bag makes it easier to extract insects from the trap and to undertake the initial processing steps in the field with minimal handling of samples, thereby reducing the risks of contamination across samples. Despite these advantages, however, it is necessary to bear in mind some limitations and potential shortcomings, which are similar to those already described for light traps in general. For instance, light traps only sample part of the nocturnal flying insect community, as many species are not attracted to light and/or have no nocturnal activity and thus cannot be represented in the samples (Young 2005). Also, the area effectively sampled by a light trap (i.e., trapping radius) may be conditional on vegetation density affecting penetration of the light source (Bowden 1982), with traps set in areas with higher visibility (e.g., grasslands) possibly attracting individuals from farther away than those set in more cluttered habitats (e.g., forests and shrublands). Therefore, when designing a study based on light traps such as ours, researchers need to be aware of these and other potential constrains, using best practices developed from previous studies to control or correct eventual sampling biases (Bowden 1982, Young 2005).

The comparisons of species composition between morphological identification and metabarcoding revealed a reasonable matching when using the ‘individuals’ and ‘legs’ bulk samples. Nevertheless, about 35% of the species occurrences were non-congruent between methods, likely due to limitations of species identification using morphology, metabarcoding, or both. A significant part of the mismatches was probably due to the difficulties of morphological identification of small microlepidoptera species, as species occurrences detected with both methods raised markedly from micro- to macrolepidoptera. Because of these difficulties, morphological examination sometimes assigned an individual to a species while metabarcoding recorded another similar species from the same genus. In a few cases, mismatches involved morphology assigning several similar individuals to the same species, while metabarcoding revealed that a few of these belonged to a second or even a third species. These results point out the limitations of using morphological examination as the benchmark for metabarcoding validation, even in cases such as ours where the fauna studied was reasonably well known (Corley 2015), and a very experienced specialist was involved in the study (M. F. V. Corley). This is in line with the results of recent efforts to DNA barcode the Portuguese moth fauna, which has revealed several new species, some of which had been previously misidentified (e.g., Corley & Ferreira 2019, Corley et al. 2019). In a few cases, mismatches were associated with limitations of metabarcoding, with the markers used being unable to discriminate between closely related species, while morphology readily provided species-level identifications. Metabarcoding also failed to identify species for which there were no DNA barcodes available at the time of the study, which is a general problem affecting metabarcoding studies. Finally, metabarcoding possibly produced a few false positives, such as *Agrotis bigramma, Leucochlaena oditis* and *Luperina Testacea*, as these species are known to fly only in late summer and early autumn while our samples were collected in July. All these species were represented by just a few reads, and possibly resulted from the assignment of reads to incorrect samples (cross-talk) (e.g., Edgar 2018), as our samples were run together with light trap samples collected in September. Overall, therefore, these comparisons suggest that metabarcoding provided an accurate description of the communities represented in the ‘individuals’ and ‘legs’ bulk samples, with most errors resulting from problems of unresolved taxonomy or incomplete databases, though there might have been errors associated with metabarcoding itself.

Comparisons between morphology and metabarcoding involving the ‘mixture’ bulk sample provided the lowest congruence levels in species occurrences. Errors were mainly due to metabarcoding detecting a large number of species not recorded through morphology, while only a very few species were detected through morphology but not metabarcoding. These results are likely a consequence of the ‘mixture’ bulks retaining many small individuals from unidentified species, particularly microlepidoptera, which contributed to increase taxonomic diversity. Furthermore, there were probably body parts retained in the mixture that could not be detected, let alone identified, and that further contributed to increase diversity. Finally, some species could have been detected through the stomach contents of carnivorous arthropods (Sheppard et al. 2005; da Silva et al. 2019), and not directly through individuals collected in the trap. Additionally, although eventual problems of lab and field contamination inflating numbers of taxa could not be totally discarded, we believe that these should have had little influence in the results, as this problem was not detected for the ‘individuals’ and ‘legs’ bulks. Therefore, it is unlikely that the higher number of taxa recorded through metabarcoding in ‘mixture’ bulks were false positives, except in the few cases involving cross talk errors. Instead, the higher number of taxa likely reflected the much-increased sensitivity of metabarcoding to estimate species diversity in relation to the conventional morphological approach. This result also suggests that metabarcoding validation studies using only artificial mock communities may be underestimating the true power of the technique to reveal the diversity of real communities, and so the use in validation of actual field samples is highly recommended.

## Conclusions

The customised light trap described in this study in combination with DNA metabarcoding offers a relatively simple and cost-effective approach to describe communities of nocturnal flying insects in large scale studies. Although like other approaches based on metabarcoding it cannot provide information on species abundances (Elbrecht & Leese 2015; Piñol et al. 2015), it offers the ability to process hundreds to thousands of samples at high taxonomic resolution in a relatively short time frame, which can hardly be achieved through conventional morphological approaches (Ji et al. 2013). A key advantage of our approach is that a large number of insect specimens can be collected at a number of sites in a single night by a small field team (Zenke et al. 2020), which can be particularly advantageous if the study is carried out in remote areas, while other sampling devices such as Malaise traps are usually more labour intensive and require deployment in the field for much longer periods (e.g., Matthews & Matthews 1971; Häuser & Riede 2015). Like any sampling device, however, our method is biased towards some species groups such as flying Lepidoptera, Diptera and Coleoptera that are attracted to light, and thus a combination of sampling methods should be used in studies requiring a comprehensive account of entire insect communities (e.g., Yang et al. 2014, Marquina et al. 2019). To further enhance the value of our approach, it would be important to increase the DNA barcode reference databases for hyper-diverse insect groups such as Diptera (e.g., Ferreira et al 2020), to apply bioinformatic procedures correcting for cross talk problems causing false positives in metabarcoding (e.g., Edgar 2018), to standardise procedures that may affect catchability of different insect groups such as for instance the light source and intensity, and the weather and moonlight conditions (Young 2005), and to further assess potential errors and limitations such as variations in effective catching distance across habitats with different vegetation cluttering (e.g., forests with or without undergrowth; Young 2005). Overall, we suggest that the methodological pipeline proposed in this study provides a convenient approach for the standardised monitoring of spatio-temporal variations in the diversity and composition of complex nocturnal insect communities, with the potential to help addressing pressing societal challenges such as global insect declines.

## References

Aizpurua, O., Budinski, I., Georgiakakis, P., Gopalakrishnan, S., Ibañez, V., Mata, V., … Alberdi, A. (2018). Agriculture shapes the trophic niche of a bat preying multiple pest arthropods across Europe: Evidence from DNA metabarcoding. Molecular Ecology, 27, 815–825. doi: 10.1111/mec.14474

Ballare, K. M., Pope, N. S., Castilla, A. R., Cusser, S., Metz, R. P., & Jha, S. (2019). Utilizing field collected insects for next generation sequencing: Effects of sampling, storage, and DNA extraction methods. Ecology and Evolution, 9, 13690–13705. doi: 10.1002/ece3.5756

Barsoum, N., Bruce, C., Forster, J., Ji, Y. Q., & Douglas, W. Y. (2019). The devil is in the detail: Metabarcoding of arthropods provides a sensitive measure of biodiversity response to forest stand composition compared with surrogate measures of biodiversity. Ecological Indicators, 101, 313–323. doi: 10.1016/j.ecolind.2019.01.023

Basset, Y., & Lamarre, G. P. A. (2019). Toward a world that values insects. Science, 364, 1230–1231. doi:1231. 10.1126/science.aaw7071

Bates, D., Mächler, M., Bolker, B., & Walker, S. (2015). Fitting linear mixed-effects models using lme4. Journal of Statistical Software, 67, 1–48. doi: 10.18637/jss.v067.i01

Bell, J. R., Blumgart, D., & Shortall, C. R. 2020. Are insects declining and at what rate? An analysis of standardised, systematic catches of aphid and moth abundances across Great Britain. Insect Conservation and Diversity, 13, 115–126. doi: 10.1111/icad.12412

Blanckenhorn, W. U., Rohner, P. T., Bernasconi, M. V., Haugstetter, J., & Buser, A. (2016). Is qualitative and quantitative metabarcoding of dung fauna biodiversity feasible?. Environmental Toxicology and Chemistry, 35, 1970–1977. doi: 10.1002/etc.3275

Brandon-Mong, G. J., Gan, H. M., Sing, K. W., Lee, P. S., Lim, P. E., & Wilson, J. J. (2015). DNA metabarcoding of insects and allies: an evaluation of primers and pipelines. Bulletin of Entomological Research, 105, 717–727. doi: 10.1017/S0007485315000681

Braukmann, T. W., Ivanova, N. V., Prosser, S. W., Elbrecht, V., Steinke, D., Ratnasingham, S., … & Hebert, P. D. (2019). Metabarcoding a diverse arthropod mock community. Molecular Ecology Resources, 19, 711–727. doi: 0.1111/1755-0998.13008

Bowden, J. (1982). An analysis of factors affecting catches of insects in light-traps. Bulletin of Entomological Research, 72, 535–556. doi: 10.1017/S0007485300008579

Bush, A., Monk, W. A., Compson, Z. G., Peters, D. L., Porter, T. M., Shokralla, S., … Baird, D. J. (2020). DNA metabarcoding reveals metacommunity dynamics in a threatened boreal wetland wilderness. Proceedings of the National Academy of Sciences, 117, 8539–8545. doi: 10.1101/819714

Corley, M F. V. (2015). Lepidoptera of Continental Portugal. Faringdon, UK: Martin Corley.

Corley, M., Ferreira, S., & Mata, V. A. (2019). *Ypsolopha rhinolophi* sp. nov. (Lepidoptera: Ypsolophidae), a new species from Portugal and France unveiled by bats. Zootaxa, 4609, 565–573. doi: 10.11646/zootaxa.4609.3.10

Corley, M., & Ferreira, S. (2019). A taxonomic revision of the Western Palaearctic genus *Cacochroa* Heinemann, 1870 (Lepidoptera, Depressariidae, Cryptolechiinae) with description of a new genus and a new species. Zootaxa, 4683, 197–214. doi: 10.11646/zootaxa.4683.2.2

deWaard, J. R., Levesque-Beaudin, V., deWaard, S. L., Ivanova, N. V., McKeown, J. T., Miskie, R., … Sones, J. E. (2019). Expedited assessment of terrestrial arthropod diversity by coupling Malaise traps with DNA barcoding. Genome, 62, 85–95. doi: 10.1139/gen-2018-0093

da Silva, L. P., Mata, V. A., Lopes, P. B., Pereira, P., Jarman, S. N., Lopes, R. J., & Beja, P. (2019). Advancing the integration of multi-marker metabarcoding data in dietary analysis of trophic generalists. Molecular Ecology Resources, 19, 1420–1432. doi: 10.1111/1755-0998.13060

Dillon, N., Austin, A. D., & Bartowsky, E. (1996). Comparison of preservation techniques for DNA extraction from hymenopterous insects. Insect Molecular Biology, 5, 21–24. doi: 10.1111/j.1365-2583.1996.tb00036.x

Douglas, W. Y., Ji, Y., Emerson, B. C., Wang, X., Ye, C., Yang, C., & Ding, Z. (2012). Biodiversity soup: metabarcoding of arthropods for rapid biodiversity assessment and biomonitoring. Methods in Ecology and Evolution, 3, 613–623. doi: 10.1111/j.2041-210X.2012.00198.x

Ebach, M. C., Valdecasas, A. G., Wheeler, Q. D. (2011). Impediments to taxonomy and users of taxonomy: accessibility and impact evaluation. Cladistics, 27, 550–557. doi: 10.1111/j.1096-0031.2011.00348.x

Elbrecht, V., & Leese, F. (2015). Can DNA-based ecosystem assessments quantify species abundance? Testing primer bias and biomass—sequence relationships with an innovative metabarcoding protocol. PloS one, 10, e0130324. doi: 10.1371/journal.pone.0130324

Elbrecht, V., Vamos, E. E., Meissner, K., Aroviita, J., & Leese, F. (2017). Assessing strengths and weaknesses of DNA metabarcoding-based macroinvertebrate identification for routine stream monitoring. Methods in Ecology and Evolution, 8, 1265–1275. doi: 10.1111/2041-210X.12789

Elbrecht, V., & Leese, F. (2017). Validation and development of COI metabarcoding primers for freshwater macroinvertebrate bioassessment. Frontiers in Environmental Science, 5, 11. doi: 10.3389/fenvs.2017.00011

Elbrecht, V., Braukmann, T. W., Ivanova, N. V., Prosser, S. W., Hajibabaei, M., Wright, M., … Steinke, D. (2019). Validation of COI metabarcoding primers for terrestrial arthropods. PeerJ, 7, e7745. doi: 10.7717/peerj.7745

Ferreira, S., Andrade, R., Gonçalves, A., Sousa, P., Paupério, J., Fonseca, N., & Beja, P. (2020). The InBIO Barcoding Initiative Database: DNA barcodes of Portuguese Diptera 01. Biodiversity Data Journal, 8, e49985. doi: 10.3897/BDJ.8.e49985

Frøslev, T. G., Kjøller, R., Bruun, H. H., Ejrnæs, R., Brunbjerg, A. K., Pietroni, C., & Hansen, A. J. (2017). Algorithm for post-clustering curation of DNA amplicon data yields reliable biodiversity estimates. Nature Communications, 8, 1–11. doi: 10.1038/s41467-017-01312-x

Gaston, K.J., 2019. Nighttime Ecology: The “Nocturnal Problem” Revisited. American Naturalis 193, 481–502. doi: 10.1086/702250

Gueuning, M., Ganser, D., Blaser, S., Albrecht, M., Knop, E., Praz, C., & Frey, J. E. (2019). Evaluating next-generation sequencing (NGS) methods for routine monitoring of wild bees: Metabarcoding, mitogenomics or NGS barcoding. Molecular Ecology Resources, 19, 847–862. doi: 10.1111/1755-0998.13013

Hallmann, C. A., Sorg, M., Jongejans, E., Siepel, H., Hofland, N., Schwan, H., … de Kron, H. (2017), More than 75 percent decline over 27 year in total flying insect biomass in protected areas. PLoS ONE 12, e0185809. doi: 10.1371/journal.pone.0185809

Häuser, C. L., & Riede, K. (2015). Field methods for inventorying insects. In M. F. Watson, C. H. C. Lyal & C. A. Pendry (Eds.), Descriptive taxonomy: the foundation of biodiversity research (pp. 190–213). Cambridge: Cambridge University Press.

Hering, D., Borja, A., Jones, J. I., Pont, D., Boets, P., Bouchez, A., … & Kelly, M. (2018). Implementation options for DNA-based identification into ecological status assessment under the European Water Framework Directive. Water Research, 138, 192–205. doi: 10.1016/j.watres.2018.03.003

Ji, Y., Ashton, L., Pedley, S. M., Edwards, D. P., Tang, Y., Nakamura, A., … & Larsen, T. H. (2013). Reliable, verifiable and efficient monitoring of biodiversity via metabarcoding. Ecology Letters, 16, 1245–1257. doi: 10.1111/ele.12162

Macgregor, C. J., Pocock, M. J., Fox, R., & Evans, D. M. (2015). Pollination by nocturnal Lepidoptera, and the effects of light pollution: a review. Ecological Entomology, 40, 187–198. doi: 10.1111/een.12174

Marquina, D., Esparza-Salas, R., Roslin, T., & Ronquist, F. (2019). Establishing arthropod community composition using metabarcoding: Surprising incosnsistencies between soil samples and preservative ethanol and homogenate from Malaise trap catches. Molecular Ecology Resources, 19, 1516–1530. doi: 10.1111/1755-0998.13071

Mata, V. A., Amorim, F., Corley, M. F.V., McCracken, G. F., Rebelo, H. & Beja, P. (2016). Female dietary bias towards large migratory moths in the European free-tailed bat (*Tadarida teniotis*). Biology Letters, 12, 20150988. doi: 10.1098/rsbl.2015.0988

Matthews, R. W., & Matthews, J. R. (1971). The Malaise trap: its utility and potential for sampling insect populations. The Great Lakes Entomologist, 4, 117–122.

Edgar, R. (2018). UNCROSS2: identification of cross-talk in 16S rRNA OTU tables. bioRxiv, 400762.

Oksanen, J., Blanchet, F. G., Friendly, M., Kindt, R., Legendre, P., McGlinn, D., … Wagner, H. (2018). vegan: Community Ecology Package. R package version 2.5-3. 2018. https://cran.r-project.org/package=vegan

Owens, A., Cochard, P., Durrant, J., Perkin, E., & Seymoure, B. (2020). light pollution is a driver of insect declines. Biological Conservation, 241: 108259. doi: 10.1016/j.biocon.2019.108259

Piñol, J., Mir, G., Gomez-Polo, P., & Agustí, N. (2015). Universal and blocking primer mismatches limit the use of high-throughput DNA sequencing for the quantitative metabarcoding of arthropods. Molecular Ecology Resources, 15, 819–830. doi: 10.1111/1755-0998.12355

Piper, A. M., Batovska, J., Cogan, N. O., Weiss, J., Cunningham, J. P., Rodoni, B. C., & Blacket, M. J. (2019). Prospects and challenges of implementing DNA metabarcoding for high-throughput insect surveillance. GigaScience, 8, giz092. doi: 10.1093/gigascience/giz092

R Core Team (2018). R: A language and environment for statistical computing.R Foundation for Statistical Computing, Vienna, Austria. URL https://www.R-project.org/

Rognes, T., Flouri, T., Nichols, B., Quince, C., & Mahé, F. (2016). VSEARCH: a versatile open source tool for metagenomics. PeerJ, 4, e2584. doi: 10.7717/peerj.2584

Sánchez-Bayo, F. & Wyckhuys. (2019). Worldwide decline of the entomofauna: A review of its drivers. Biological Conservation, 232, 8–27. doi: 10.1016/j.biocon.2019.01.020

Sheppard, S. K., Bell, J., Sunderland, K. D., Fenlon, J., Skervin, D., & Symondson, W. O. C. (2005). Detection of secondary predation by PCR analyses of the gut contents of invertebrate generalist predators. Molecular Ecology, 14(14), 4461–4468. doi: 10.1111/j.1365-294X.2005.02742.x

Sierro, A., Arlettaz, R., Naef-Daenzer, B., Strebel, S., & Zbinden, N. (2001). Habitat use and foraging ecology of the nightjar (*Caprimulgus europaeus*) in the Swiss Alps: towards a conservation scheme. Biological conservation, 98, 325–331. doi: 10.1016/S0006-3207(00)00175-0

Wagner, D. L. (2020). Insect declines in the Anthropocene. Annual Review of Entomology, 65, 457–480. doi: 10.1146/annurev-ento-011019-025151

Yang, C., Wang, X., Miller, J. A., de Blécourt, M., Ji, Y., Yang, C., … & Yu, D. W. (2014). Using metabarcoding to ask if easily collected soil and leaf-litter samples can be used as a general biodiversity indicator. Ecological Indicators, 46, 379–389. doi: 10.1016/j.ecolind.2014.06.028

Young, M. (2005). Insects in flight. In S. Leather (Ed.), Insect sampling in forest ecosystems (pp. 116–145). Malden, MS: Blackwell Science.

Zenker, M. A., Specht, A. & Fonseca, V. G. (2020). Assessing insect biodiversity with automatic light in Brazil: Pearls and pitfalls of metabarcoding samples in preservative ethanol. Ecology and Evolution, 10, 2352–2366. doi: 10.1002/ece3.6042

