## Supplementary Methods for "Efficient assessment of nocturnal flying insect communities by combining automatic light traps and DNA metabarcoding"

**Arthropod DNA extraction from bulk samples using the E.Z.N.A.® Tissue DNA Kit**

**Before starting:**

- Set the oven at 56ºC and place custom lysis washing buffer (0.1 M Tris–HCl, 0.1 M EDTA, 0.01 M NaCl, 1% N-lauroylsarcosine, pH 7.5-8) and E.Z.N.A. elution buffer to incubate;
- Set the dry-bath to 70ºC
- Clean the working space and material with bleach and water and leave it under the UV-light for at least 15 mins;
- Use filter tips at all steps;
- Use a negative control for each batch.

**Procedure:**

1. Prepare the number of 2mL tubes needed for the extraction and weight 80 mg – 100 mg of homogenized insect powder.
2. Distribute 1000 μL of custom lysis washing buffer and 25 μL of OB Protease per tube. Carefully, dispense buffer to tube in order to avoid disturbing the powder and consequently cross contaminate samples.
3. Quickly vortex tubes and incubate them in the oven for 30 minutes.
4. While samples are incubating, prepare the following per number of extractions:

- one 2mL tube with half Inhibitex tablet;
- one 2ml tube with 25 μL of OB Protease

5. Short-spin tubes and transfer up to 700 μL of the supernatant to the tube with Inhibitex.
6. Vortex for 1 min and centrifuge for 30 s at 14000rpm.
7. Transfer up to 500 μL of the supernatant (avoid disturbing the pellet) to the 2mL tube containing OB Protease.
8. Add 200 μL of BL buffer. Vortex at maximum speed for 15 seconds and short-spin.
9. Place the tubes in the dry-bath for 10 minutes.
10. Short-spin the tubes and add 400 μL of ethanol 100% and vortex at maximum speed for 20 seconds. Short-spin samples.

***Safe point to stop!*** *If needed, you can put samples in the freezer (covered in aluminum foil to avoid contaminations). If stopping, when re-starting the protocol allow samples to come to room temperature for a few minutes.*

***If using the centrifuge see steps 11 to 18. If using QIAGEN Vacuum Pump see steps 19to 30.***

**When using the centrifuge:**

11. Transfer up to 600 µl of supernatant to the column with a collector tube. Centrifuge at 1000rpm for 1 minute.
12. Place the column in a new collection tube.
13. Repeat step 16 and 17 if you still have volume left from step 15.
14. Place the column in a new collection tube. Add 500 µl of HB Buffer. Centrifuge at 1000rpm for 1 minute.
15. Place the column in a new collection tube. Add 700 µl of DNA Wash Buffer. Centrifuge at 1000rpm for 1 minute.
16. Repeat previous step.
17. Place the column in a new collection tube. Centrifuge ate 14000rpm for 2 minutes to completely dry the membrane.
18. Proceed to step 26.

**When using the QIAvac:**

*Do not forget to place a VacConnector between the QIAvac and EZNA column.*

19. Transfer up to 600 µl of supernatant to the column placed in the QIAvac. Repeat if necessary. Carefully, so samples do not overflow.
20. Turn on the Vacuum pump. The ideal vacuum pressure should be between -80 and -90 kPa.
21. Add 500 µl of HB Buffer. If one or more columns appears to be clogged turn off the pump when possible. Place the clogged column in a collector tube and centrifuge at 1000rpm for 1 minute. Return the column to its position in the QIAvac.
22. Add 700 µl of DNA Wash Buffer.
23. Repeat previous step.
24. Allow the columns membrane to completely dry for 5 up to 10 mins (depends on the room temperature).
25. Proceed to step 26.

**DNA Elution:**

26. Transfer the column plate to 1.5mL labelled tube. Add 50 µL Elution Buffer and Incubate at room temperature for 5 minutes. Centrifuge at 10000 rpm for 1 minutes. Repeat, for a second elution if desired.
