## Supplementary Table S1 for "Efficient assessment of nocturnal flying insect communities by combining automatic light traps and DNA metabarcoding"

**SUPPLEMENTARY TABLES**

**Table S1.** Species of Lepidoptera captured in 12 light traps set in cork oak woodlands in NE Portugal (July 2017) and detected with either metabarcoding (Meta) or morphology (Morpho). Comparisons between metabarcoding and morphology were made using bulks produced using either one individual per species (‘individuals’ bulk), one leg per species (‘legs’ bulk), or the unsorted mixture of individuals except those used in the previous two bulks (‘mixture’ bulk). We provide the number of samples in which a species was detected, for each bulk type and method.

| **Species** | **‘individuals' bulk** | | **‘legs' bulk** | | **‘mixture' bulk** | |
| --- | --- | --- | --- | --- | --- | --- |
|  | **Meta** | **Morpho** | **Meta** | **Morpho** | **Meta** | **Morpho** |
| **Autostichidae** |  |  |  |  |  |  |
| *Apatema apolausticum* |  |  |  |  | 5 |  |
| *Apatema mediopallidum* |  | 1 |  | 1 |  |  |
| *Apatema sp. 1* | 1 |  | 1 |  |  |  |
| *Apatema sp. 2* |  |  |  |  | 4 |  |
| *Oegoconia quadripuncta* |  |  |  |  | 2 |  |
| *Symmoca alhambrella* |  |  |  |  | 2 |  |
| *Symmoca revoluta* |  |  |  |  | 1 |  |
| *Symmoca signatella* | 3 | 3 | 2 | 3 | 8 |  |
| *Symmoca* sp. 1 | 1 |  | 1 |  | 5 |  |
| **Brachodidae** |  |  |  |  |  |  |
| *Brachodes funebris* | 6 |  | 6 |  | 7 |  |
| *Brachodes gaditana* | 3 | 7 | 2 | 7 | 5 | 4 |
| **Coleophoridae** |  |  |  |  |  |  |
| *Coleophora struella* |  |  |  |  | 2 |  |
| **Cosmopterigidae** |  |  |  |  |  |  |
| *Eteobalea intermediella* |  |  |  |  | 2 |  |
| *Pyroderces argyrogrammos* |  |  |  |  | 1 |  |
| *Vulcaniella grabowiella* |  |  |  |  | 2 |  |
| **Crambidae** |  |  |  |  |  |  |
| *Agrotera nemoralis* | 1 | 1 | 1 | 1 |  |  |
| *Anarpia incertalis* | 2 | 2 | 2 | 2 | 1 | 2 |
| *Catoptria pinella* | 2 | 2 | 2 | 2 |  |  |
| *Eudonia mercurella* | 10 | 11 | 10 | 11 | 10 | 10 |
| *Mecyna asinalis* |  |  |  |  | 1 |  |
| *Metasia cuencalis* | 3 | 1 | 3 | 1 | 8 |  |
| *Patania crocealis* | 1 | 1 | 1 | 1 |  |  |
| *Pyrausta aurata* |  |  |  |  | 1 |  |
| **Depressariidae** |  |  |  |  |  |  |
| *Agonopterix atomella* | 1 |  | 1 |  |  |  |
| *Agonopterix scopariella* | 3 | 3 | 3 | 3 |  |  |
| *Depressaria* sp. 1 |  | 1 |  | 1 |  |  |
| *Depressaria adustatella* | 1 |  | 1 |  | 1 |  |
| **Drepanidae** |  |  |  |  |  |  |
| *Watsonalla uncinula* | 5 | 6 | 5 | 6 | 7 | 6 |
| **Epermeniidae** |  |  |  |  |  |  |
| *Epermenia aequidentellus* |  |  |  |  | 1 |  |
| **Erebidae** |  |  |  |  |  |  |
| *Catephia alchymista* | 1 | 1 | 1 | 1 | 2 |  |
| *Catocala nymphagoga* | 10 | 10 | 10 | 10 | 10 | 8 |
| *Dysgonia algira* | 1 | 1 | 1 | 1 | 1 |  |
| *Eilema uniola* | 1 |  |  |  | 1 |  |
| *Eublemma candidana* | 1 |  | 1 |  |  |  |
| *Eublemma parva* | 6 | 6 | 5 | 6 | 2 | 2 |
| *Eublemma pura* | 4 | 5 | 4 | 5 | 1 |  |
| *Lymantria dispar* | 12 | 12 | 12 | 12 | 11 | 9 |
| *Ocneria rubea* | 1 | 1 | 1 | 1 | 1 |  |
| *Phragmatobia fuliginosa* | 1 | 1 | 1 | 1 |  |  |
| *Zethes insularis* | 1 | 1 | 1 | 1 |  |  |
| **Gelechiidae** |  |  |  |  |  |  |
| *Anacampsis scintillella* | 2 |  | 2 |  | 1 |  |
| *Anacampsis timidella* |  |  |  |  | 2 |  |
| *Aproaerema anthyllidella* | 1 | 1 | 1 | 1 | 2 |  |
| *Aproerema polychromella* |  | 1 |  | 1 | 2 |  |
| *Aproaerema* sp. 1 | 1 |  | 1 |  |  |  |
| *Aproaerema sp. 2* |  |  |  |  | 3 |  |
| *Aristotelia decoratella* |  |  |  |  | 2 |  |
| *Aristotelia ericinella* | 1 | 2 | 1 | 2 | 5 |  |
| *Aroga pascuicola* |  |  |  |  | 1 |  |
| *Bryotropha senectella* |  |  |  |  | 1 |  |
| *Bryotropha vondermuhlli* |  |  |  |  | 5 |  |
| *Carpatolechia decorella* |  |  |  |  | 3 |  |
| *Epidola stigma* | 2 | 2 | 2 | 2 | 7 |  |
| *Gelechiidae* sp. 1 |  |  |  |  | 1 |  |
| *Gladiovalva badidorsella* | 1 |  | 1 |  |  |  |
| *Metzneria aestivella* | 1 |  |  |  | 2 |  |
| *Neofriseria hitadoella* | 6 | 7 | 6 | 7 | 9 | 3 |
| *Neofriseria singula* |  |  |  |  | 1 |  |
| *Neotelphusa cisti* | 1 |  | 2 |  | 2 |  |
| *Neotelphusa* sp. 1 |  | 1 |  | 1 |  |  |
| *Pseudotelphusa occidentella* | 2 | 1 | 1 | 1 | 5 |  |
| *Stomopteryx flavipalpella* |  |  |  |  | 2 |  |
| *Stomopteryx lusitaniella* |  |  |  |  | 5 |  |
| *Stomopteryx remissella* |  |  |  |  | 2 |  |
| *Teleiopsis diffinis* |  |  |  |  | 2 |  |
| *Xenolechia aethiops* |  |  |  |  | 1 |  |
| **Geometridae** |  |  |  |  |  |  |
| *Brachyglossina hispanaria* | 1 | 1 | 1 | 1 | 4 |  |
| *Camptogramma bilineata* | 3 | 3 | 3 | 3 | 1 | 1 |
| *Charissa mucidaria* | 1 | 1 | 1 | 1 | 1 |  |
| *Crocallis albarracina/elinguaria* |  |  |  |  | 1 |  |
| *Cyclophora puppillaria* | 4 | 4 | 4 | 4 | 5 | 3 |
| *Cyclophora serveti* |  |  |  |  | 1 |  |
| *Gymnoscelis rufifasciata* | 2 | 2 | 2 | 2 | 3 |  |
| *Idaea belemiata* |  | 12 |  | 12 |  | 11 |
| *Idaea circuitaria* | 2 | 2 | 2 | 2 | 2 |  |
| *Idaea consanguiberica* | 1 |  | 1 |  | 7 |  |
| *Idaea eugeniata* | 1 | 1 | 1 | 1 | 6 |  |
| *Idaea exilaria* | 2 | 2 | 2 | 2 | 3 |  |
| *Idaea infirmaria* | 7 | 9 | 8 | 9 | 10 | 7 |
| *Idaea mustelata* | 7 | 7 | 7 | 7 | 11 | 5 |
| *Idaea nigrolineata* |  | 1 |  | 1 |  |  |
| *Idaea obsoletaria* | 9 |  | 8 |  | 12 |  |
| *Idaea rhodogrammaria* | 7 | 4 | 6 | 4 | 10 | 3 |
| *Idaea* sp. |  | 1 |  | 1 |  |  |
| *Idaea subsericeata* |  |  |  |  | 3 |  |
| *Menophra abruptaria* | 3 | 2 | 3 | 2 |  |  |
| *Menophra japygiaria* | 1 | 2 | 2 | 2 |  |  |
| *Nychiodes andalusiaria* | 8 | 8 | 8 | 8 | 5 | 3 |
| *Pachycnemia hippocastanaria* |  |  |  |  | 2 |  |
| *Phaiogramma etruscaria* | 2 | 2 | 2 | 2 | 2 |  |
| *Plagodis dolabraria* | 1 | 1 | 1 | 1 | 2 |  |
| *Pseudoterpna coronillaria* | 5 | 4 | 5 | 4 | 6 | 2 |
| *Rhoptria asperaria* | 5 | 5 | 5 | 5 | 3 | 1 |
| *Scopula marginepunctata* | 1 |  | 1 |  | 1 |  |
| *Scopula submutata* |  | 1 |  | 1 |  |  |
| *Tephronia sepiaria* | 3 | 4 | 3 | 4 | 2 |  |
| **Gracillariidae** |  |  |  |  |  |  |
| *Phyllonorycter endryella* |  |  |  |  | 2 |  |
| *Phyllonorycter millierella* |  |  |  |  | 1 |  |
| *Phyllonorycter roboris* |  |  |  |  | 1 |  |
| *Phyllonorycter trifasciella* |  |  |  |  | 1 |  |
| *Triberta helianthemella* | 1 |  |  |  | 5 |  |
| **Lasiocampidae** |  |  |  |  |  |  |
| *Phyllodesma suberifolia* | 9 | 7 | 8 | 7 | 5 | 4 |
| *Psilogaster loti* | 5 | 4 | 4 | 4 | 5 | 1 |
| **Lecithoceridae** |  |  |  |  |  |  |
| *Eurodachtha canigella* |  | 3 |  | 3 |  | 2 |
| *Eurodachtha siculella/canigella* | 1 |  | 1 |  | 7 |  |
| *Lecithoceridae* 1 | 1 |  | 1 |  | 7 |  |
| **Momphidae** |  |  |  |  |  |  |
| *Mompha miscella* |  | 1 |  | 1 |  |  |
| *Mompha subbistrigella* |  |  |  |  | 2 |  |
| **Nepticulidae** |  |  |  |  |  |  |
| *Stigmella eberhardi* |  |  |  |  | 3 |  |
| *Zimmermannia hispanica* |  |  |  |  | 3 |  |
| *Zimmermannia* sp. 1 |  |  |  |  | 2 |  |
| **Noctuidae** |  |  |  |  |  |  |
| *Acontia lucida* | 1 | 1 | 1 | 1 |  |  |
| *Agrotis bigramma* | 3 |  |  |  | 2 |  |
| *Bryophila ravula* | 1 |  | 1 |  | 2 |  |
| *Calophasia sp.* |  | 1 |  | 1 |  |  |
| *Caradrina aspersa* | 8 | 8 | 8 | 8 | 9 | 6 |
| *Chloantha hyperici* | 1 |  | 1 |  |  |  |
| *Colocasia coryli* |  |  |  |  | 1 |  |
| *Cryphia algae* | 5 | 6 | 6 | 6 | 7 | 6 |
| *Cryphia pallida* | 2 |  | 1 |  |  |  |
| *Epilecta linogrisea* | 6 | 6 | 6 | 6 | 6 | 3 |
| *Euxoa tritici* | 1 | 1 | 1 | 1 | 1 | 1 |
| *Hecatera dysodea* | 1 | 1 | 1 | 1 | 1 |  |
| *Heliothis peltigera* |  | 1 |  | 1 |  |  |
| *Leucochlaena oditis* | 1 |  |  |  |  |  |
| *Lophoterges millierei* | 1 | 1 | 1 | 1 |  |  |
| *Luperina testacea* |  |  |  |  | 1 |  |
| *Noctua comes* | 6 | 5 | 7 | 5 | 6 | 1 |
| *Noctua fimbriata* |  | 1 |  | 1 |  | 1 |
| *Noctua janthe* | 4 | 5 | 4 | 5 | 2 |  |
| *Noctua orbona* |  | 1 |  | 1 |  | 1 |
| *Noctua tirrenica* | 1 |  | 1 |  | 2 |  |
| *Nyctobrya muralis* | 7 | 7 | 7 | 7 | 5 | 4 |
| *Paucgraphia erythrina* | 1 | 1 | 1 | 1 | 1 |  |
| *Polyphaenis sericata* | 1 | 3 | 1 | 3 | 4 |  |
| *Tyta luctuosa* | 1 | 1 | 1 | 1 | 1 |  |
| **Nolidae** |  |  |  |  |  |  |
| Meganola strigula | 3 | 4 | 3 | 4 |  |  |
| Nola tutulella |  |  |  |  | 1 |  |
| **Notodontidae** |  |  |  |  |  |  |
| *Harpyia milhauseri* | 3 | 3 | 3 | 3 | 3 |  |
| *Spatalia argentina* | 1 | 1 | 1 | 1 | 1 |  |
| *Thaumetopoea pityocampa* | 1 | 1 | 2 | 1 |  |  |
| **Nymphalidae** |  |  |  |  |  |  |
| *Maniola jurtina* | 1 | 1 | 1 | 1 | 1 |  |
| **Oecophoridae** |  |  |  |  |  |  |
| *Epicallima mercedella* | 4 | 5 | 5 | 5 | 12 | 3 |
| *Goidanichiana jourdheuillella* |  | 1 | 1 | 1 |  |  |
| *Pleurota andalusica* |  |  |  |  | 3 |  |
| *Pleurota honorella* | *1* | 1 | 1 | 1 | 1 |  |
| **Psychidae** |  |  |  |  |  |  |
| Eumasia parietariella |  |  |  |  | 1 |  |
| **Pterophoridae** |  |  |  |  |  |  |
| *Crombrugghia laetus* | 1 | 1 | 1 | 1 | 3 |  |
| *Stangeia siceliota* | 1 | 1 | 1 | 1 | 1 |  |
| **Pyralidae** |  |  |  |  |  |  |
| *Acrobasis consociella* |  |  |  |  | 1 |  |
| *Acrobasis glaucella* | 6 |  | 6 |  | 6 |  |
| *Acrobasis sodalella* |  | 2 |  | 2 |  | 1 |
| *Acrobasis sodalella/glaucella* |  | 2 |  | 2 |  | 2 |
| *Acrobasis* sp. |  | 2 |  | 2 |  | 1 |
| *Acrobasis tumidana* | 1 | 1 | 1 | 1 |  |  |
| *Aglossa brabanti* |  |  |  |  | 1 |  |
| *Ancylosis cinnamomella* | 1 | 2 | 1 | 2 | 4 |  |
| *Ancylosis oblitella* |  |  |  |  | 2 |  |
| *Ancylosis* sp. 1 | 1 |  | 1 |  |  |  |
| *Asalebria florella* | 1 |  |  |  | 3 |  |
| *Bostra obsoletalis* | 5 | 7 | 4 | 7 | 7 | 3 |
| *Cadra figulilella* |  |  |  |  | 1 |  |
| *Cadra furcatella* |  |  |  |  | 1 |  |
| *Elegia cf fallaximima* |  | 1 |  | 1 |  |  |
| *Elegia fallax* | 1 |  | 1 |  |  |  |
| *Endotricha flammealis* | 1 | 1 | 1 | 1 | 2 |  |
| *Ephestia disparella* |  |  |  |  | 2 |  |
| *Ephestia parasitella/woodiella* |  |  |  |  | 11 |  |
| *Epischnia illotella* |  |  |  |  | 1 |  |
| *Etiella zinckenella* | 6 | 6 | 6 | 6 | 3 | 1 |
| *Euzopherodes vapidella* |  |  |  |  | 1 |  |
| *Galleria mellonella* | 2 | 1 | 1 | 1 | 1 |  |
| *Homoeosoma nimbella* |  |  |  |  | 3 |  |
| *Homoeosoma sinuella* | 1 | 2 | 2 | 2 | 2 |  |
| *Lamoria anella* |  |  |  |  | 2 |  |
| *Oxybia transversella* | 1 | 3 | 2 | 3 | 4 |  |
| *Pempelia palumbella* | 6 | 7 | 7 | 7 | 5 | 4 |
| *Pempeliella ardosiella* |  |  |  |  | 3 |  |
| *Phycita roborella* | 4 | 4 | 4 | 4 | 4 | 2 |
| *Phycitodes lacteella* |  |  |  |  | 1 |  |
| *Phycitodes saxicola* |  |  |  |  | 2 |  |
| Pyralidae 1 | 1 |  | 1 |  |  |  |
| *Synaphe punctalis* | 5 | 5 | 5 | 5 | 4 | 2 |
| **Riodinidae** |  |  |  |  |  |  |
| *Reisserita zernyi* |  |  |  |  | 1 |  |
| **Scythrididae** |  |  |  |  |  |  |
| *Enolmis acanthella* |  |  |  |  | 2 |  |
| *Scythris cistorum* |  |  |  |  | 1 |  |
| *Scythris parafuscoaenea* | *2* | 4 | 3 | 4 | 5 | 1 |
| **Tineidae** |  |  |  |  |  |  |
| *Anomalotinea liguriella* | 2 | 2 | 2 | 2 | 11 | 1 |
| *Crassicornella agenjoi* |  |  |  |  | 3 |  |
| *Gaedikeia kokkariensis* |  |  |  |  | 2 |  |
| *Infurcitinea atrifasciella* |  |  |  |  | 4 |  |
| *Nemapogon agenjoi* | 2 | 2 | 2 | 2 | 7 |  |
| *Nemapogon nevadella* |  |  |  |  | 2 |  |
| *Neurothaumasia* sp. 1 |  |  |  |  | 4 |  |
| *Tenaga rhenania* |  |  |  |  | 4 |  |
| **Tischeriidae** |  |  |  |  |  |  |
| *Coptotriche angusticollella* |  |  |  |  | 1 |  |
| *Coptotriche marginea* |  |  |  |  | 2 |  |
| **Tortricidae** |  |  |  |  |  |  |
| *Bactra venosana* |  |  |  |  | 1 |  |
| *Clepsis siciliana* |  |  |  |  | 2 |  |
| *Cydia fagiglandana* | 12 | 12 | 12 | 12 | 12 | 12 |
| **Unknown** |  |  |  |  |  |  |
| Lepidoptera 1 |  |  |  |  | 1 |  |
| Lepidoptera 2 |  |  |  |  | 1 |  |
| Lepidoptera 3 |  |  |  |  | 1 |  |
| **Yponomeutidae** |  |  |  |  |  |  |
| *Paradoxus osyridellus* |  |  |  |  | 7 |  |
| **Ypsolophidae** |  |  |  |  |  |  |
| *Ypsolopha persicella* |  | 1 |  | 1 | 1 |  |
| *Ypsolopha rhinolophi* |  |  |  |  | 1 |  |
| **Total species** | 111 | 102 | 106 | 102 | 164 | 40 |
